# Overexpression of mitochondrial fusion genes enhances resilience and extends longevity

**DOI:** 10.1101/2023.10.24.563703

**Authors:** Annika Traa, Allison Keil, Zenith Rudich, Shusen Zhu, Jeremy M. Van Raamsdonk

## Abstract

The dynamicity of the mitochondrial network is crucial for meeting the ever-changing metabolic and energy needs of the cell. Mitochondrial fission promotes the degradation and distribution of mitochondria, while mitochondrial fusion maintains mitochondrial function through the complementation of mitochondrial components. Previously, we have reported that mitochondrial networks are tubular, interconnected and well-organized in young, healthy *C. elegans*, but become fragmented and disorganized with advancing age and in models of age-associated neurodegenerative disease. In this work, we examine the effects of increasing mitochondrial fission or mitochondrial fusion capacity by ubiquitously overexpressing the mitochondrial fission gene *drp-1* or the mitochondrial fusion genes *fzo-1* and *eat-3*, individually or in combination. We then measured mitochondrial function, mitochondrial network morphology, physiologic rates, stress resistance and lifespan. Surprisingly, we found that overexpression of either mitochondrial fission or fusion machinery both resulted in an increase in mitochondrial fragmentation. Similarly, both mitochondrial fission and mitochondrial fusion overexpression strains have extended lifespans and increased stress resistance, which appears to be at least partially due to the upregulation of multiple stress response pathways in these strains. Overall, our work demonstrates that increasing the expression of mitochondrial fission or fusion genes extends lifespan and improves biological resilience without promoting the maintenance of a youthful mitochondrial network morphology. This work highlights the importance of the mitochondria for both resilience and longevity.

## Introduction

While mitochondria have well-established roles in cellular metabolism and energy production, mitochondria also contribute to other crucial processes in the cell including calcium homeostasis, redox signalling, autophagy, innate immunity and programmed cell death [1]. The importance of mitochondria is further highlighted by how defects in multiple mitochondrial processes such as mitochondrial gene expression, redox homeostasis, respiratory chain assembly or membrane structure can cause inherited metabolic disorders, and contribute to age-related diseases such as neurodegeneration, diabetes and cancer [2].

The dynamicity of the mitochondrial network, where single mitochondrion connect and separate from an interconnected web of mitochondria, is governed by the fusion and fission of the inner and outer mitochondrial membranes [3]. During mitochondrial fusion in *C. elegans*, the inner mitochondrial membrane is fused by the Opa-1 homolog, EAT-3, while the outer mitochondrial membrane is fused by the mitofusin homolog, FZO-1. Mitochondrial fission occurs at ER contact sites, where ER tubules begin to constrict the mitochondria and a host of proteins, including FIS-1, FIS-2, MFF-1 and MFF-2 in *C. elegans*, recruit DRP-1 to the mitochondrial constriction site [4–8]. DRP-1 oligomerizes into a ring structure around the mitochondrial constriction site and completes the scission of both the inner and outer mitochondrial membrane using the energy produced by its hydrolysis of GTP [9–12].

Tight regulation of mitochondrial fission and fusion allows the mitochondrial network to dynamically respond to changing cellular needs. For example, conditions demanding a high energy output can be adapted to by promoting mitochondrial fusion, which allows for complementation of mitochondrial components and improved mitochondrial function [13, 14]. Alternatively, mitochondria may become dysfunctional under conditions of stress, in which case mitochondrial fission is important to facilitate the clearing of defective components by mitophagy [15, 16].

The role of mitochondrial fission and fusion during the aging process remains incompletely understood. As in other model organisms, the mitochondrial networks of *C. elegans* become fragmented and disorganized with age or in models of neurodegenerative diseases [17–26]. While the fragmentation of the mitochondrial network is presumed to be due to mitochondrial fission, large aggregates of swollen mitochondria frequently occur and are thought to be a product of overactive fusion and dysfunctional mitophagy [27]. However, despite age-associated increases in mitochondrial fragmentation, fission proteins DRP-1 and FIS1 are downregulated in aged mice and aged human endothelial cells [28–31]. In Drosophila, increased expression of Drp1 in midlife extends lifespan and health span, and aged flies display improved mitophagy and mitochondrial function [32]. Therefore, mitochondrial fission may be beneficial for healthy aging. In *C. elegans*, inhibition of the mitochondrial fusion gene *fzo-1* is reported to have no effect on lifespan, while inhibition of *eat-3* may increase lifespan [33, 34]. Furthermore, we previously reported that disruption of either *fzo-1* or *eat-3* activates multiple stress response pathways that are known to be tightly linked with mechanisms of longevity extension, including the DAF-16-mediated stress response, the SKN-1-mediated oxidative stress response and the ATFS-1-mediated mitochondrial unfolded protein response [17, 35, 36].

Paradoxically, disruption of mitochondrial fission can also promote lifespan extension. Inhibition of Drp1 is neuroprotective in mouse models of neurodegeneration [37–39]. In yeast, loss of the mitochondrial fission protein Dnm1p decreases mitochondrial fission, causes a tubular elongated mitochondrial network, and delays aging phenotypes [40]. In *C. elegans*, disrupting *drp-1* in wild-type animals has little or no effect on lifespan but disrupting *drp-1* in a neuronal model of polyglutamine toxicity improves both lifespan and health span, and decreases mitochondrial fragmentation [19]. The benefits of *drp-1* disruption may be tissue-specific as disruption of *drp-1* in a body wall muscle model of polyglutamine toxicity worsened lifespan, health span and mitochondrial network morphology [18]. Notably, disrupting *drp-1* in already long-lived *C. elegans* mutants, such as the insulin signaling mutants *daf-2* and *age-1*, drastically extends lifespan [41].

Other data suggests that promoting a balance between mitochondrial fission and fusion by disrupting both *drp-1* and *fzo-1* can extend lifespan and generates a mitochondrial network that does not become fragmented with age but instead remains elongated [33]. Furthermore, increased mitochondrial fusion is seen in multiple long-lived mutants and is required for their extended lifespan, including *daf-2* insulin signaling mutants, feeding defective *eat-2* mutants, germlineless *glp-1* mutants, and mildly impaired mitochondrial function mutant *clk-1* [42]. Together, these data suggest that decreasing mitochondrial fragmentation, or promoting mitochondrial network elongation can increase lifespan but that a balance between mitochondrial fission and fusion components may need to remain.

In this work, we evaluated how ubiquitous overexpression of mitochondrial fission and fusion machinery affects mitochondrial network morphology, animal physiology, lifespan and resistance to stress. We hypothesized that while animals with increased expression of mitochondrial fusion genes would have elongated mitochondrial networks and increased lifespans, overexpression of both fission and fusion machinery would give animals an increased capacity to perform both fission and fusion, maintain a dynamic mitochondrial network and thus best respond to stress and cellular needs. Surprisingly, we found that overexpression of either mitochondrial fission or fusion genes individually extended lifespan and increased stress resistance despite generating a fragmented mitochondrial network. Additionally, while animals with overexpression of both fission and fusion machinery did have enhanced longevity and stress resistance compared to wild-type animals, these animals exhibit decreased longevity and stress resistance compared to animals overexpressing either *drp-1, fzo-1* or *eat-3* individually. Thus, our findings suggest that overexpression of either a single mitochondrial fission or fusion gene can promote increased longevity and stress resistance, likely through the activation of key cellular stress response pathways, but that overexpression of multiple fission or fusion genes does not enhance this effect.

## Results

### Overexpression of mitochondrial fission or fusion genes causes mitochondrial fragmentation

To determine how overexpression of mitochondrial fission or fusion genes would affect mitochondrial network morphology, we expressed the mitochondrial fusion genes *eat-3* and *fzo-1* and the mitochondrial fission gene *drp-1* using ubiquitous promoters (*pro-1, rpl-28* and *eft-3* respectively). Additionally, we expressed a fluorescent reporter from tissue-specific promoters such that each overexpression (OE) strain had its own marker to facilitate crossing. Thus, *drp-1* OE was identified by fluorescence in the muscle, *fzo-1* OE by fluorescence in the pharynx, and *eat-3* OE by fluorescence in the intestine (**Figure 1A**).

**Figure 1.**
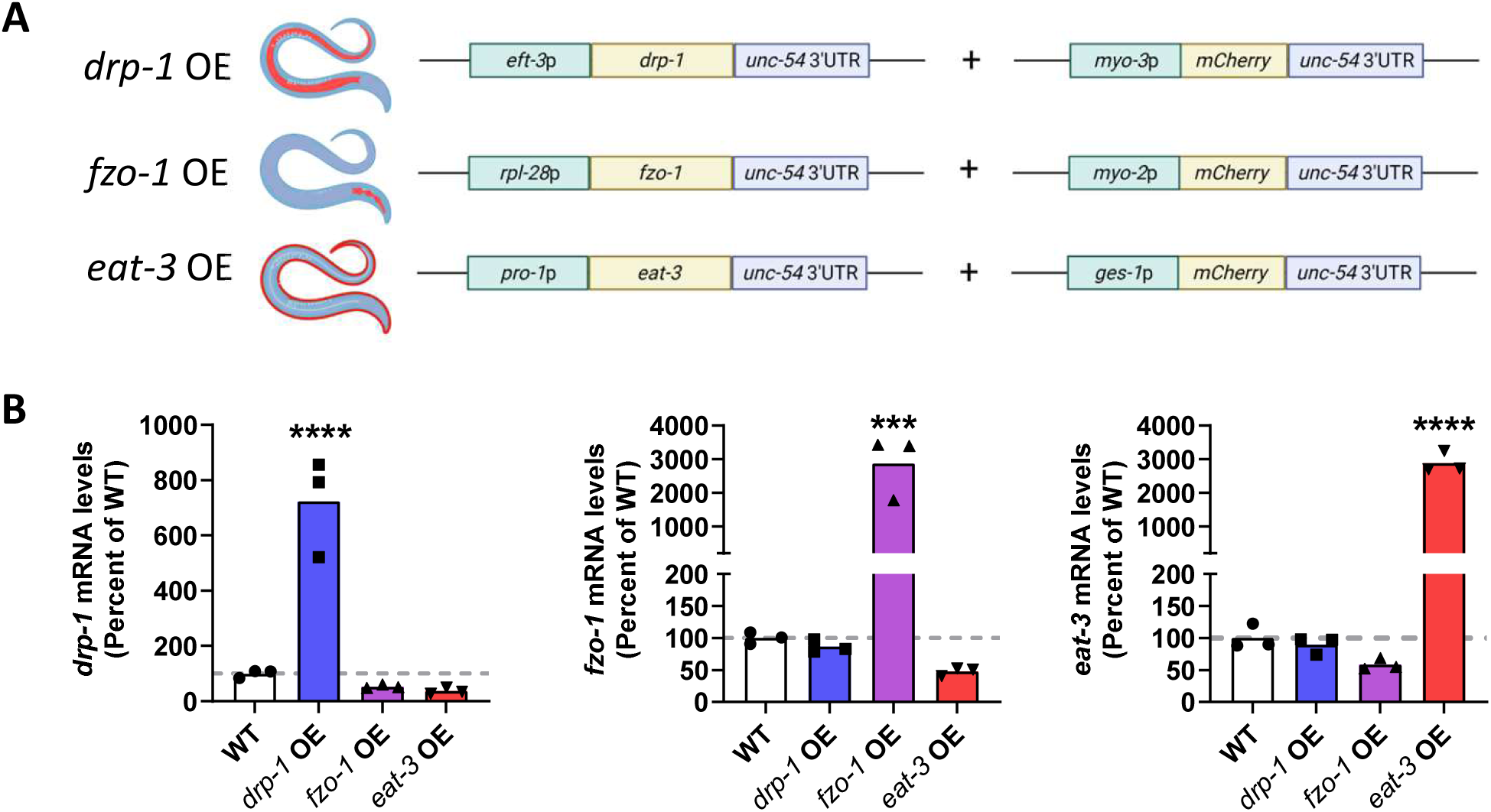
Overexpression of mitochondrial fission or fusion genes in *C. elegans.* *C. elegans* strains that overexpress mitochondrial fission or fusion genes were generated by microinjection and integration of the extrachromosomal array. (**A**) Diagram of constructs used to generate overexpression strains. The mitochondrial fission and fusion genes *drp-1, fzo-1* and *eat-3* were overexpressed under the ubiquitous promoters *eft-3, rpl-28* and *pro-1*, respectively (indicated by blue colouring in worm diagram). Each strain also expressed a red fluorescent co-injection marker in intestine (*drp-1* OE), pharynx (*fzo-1* OE) or body wall muscle (*eat-3* OE). (**B**) Quantitative RT-PCR confirmed that each overexpression strain exhibits increased expression of the intended mitochondrial fission or fusion gene. Three biological replicates were performed. Statistical significance was assessed using a one-way ANOVA with Dunnett’s multiple comparisons test. OE = overexpression. ***p<0.001, ****p<0.0001.

To validate whether the created strains had a significant increase in expression of their respective overexpressed genes, quantitative RT-PCR was used to measure the transcript levels of *drp-1, fzo-1* and *eat-3*. We found that each strain had increased mRNA levels for their corresponding gene and that overexpression of the mitochondrial fusion genes was nearly 4 times higher than overexpression of the mitochondrial fission gene *drp-1* (**Figure 1B**).

The mitochondrial network morphology of the overexpression strains was evaluated by crossing each strain with animals expressing mitochondrially-targeted GFP in body wall muscle cells. At day 1 of adulthood, overexpression of *drp-1, fzo-1*, or *eat-3* significantly increased mitochondrial fragmentation as evaluated by mitochondrial number, area, circularity and length (**Figure 2A,B**).

**Figure 2.**
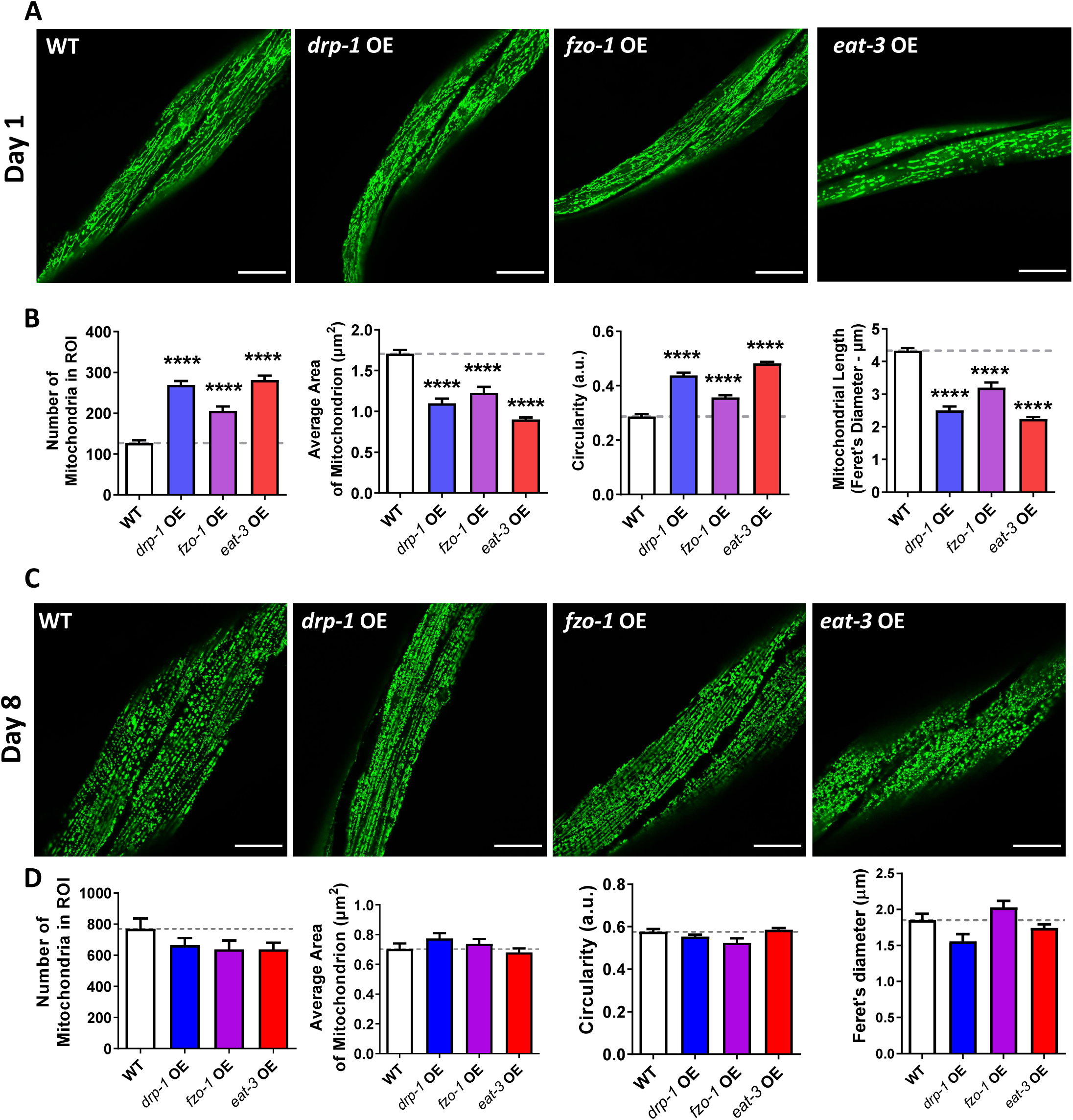
Overexpression of mitochondrial fission or fusion genes causes mitochondrial fragmentation. The mitochondrial morphology resulting from overexpression of mitochondrial fission and fusion genes was assessed at day 1 and day 8 of adulthood. (**A**) At day 1 of adulthood, overexpression of *drp-1, fzo-1* or *eat-3* resulted in mitochondrial fragmentation. (**B**) Quantification of mitochondrial morphology showed that these worms have an increased number of mitochondria, decreased mitochondrial area, increased circularity and decreased length, all consistent with increased fragmentation. (**C**) At day 8 of adulthood, mitochondrial morphology in the strains overexpressing *drp-1, fzo-1* or *eat-3* was similar to that in wild-type worms. (**D**) Quantification of mitochondrial morphology at day 8 of adulthood revealed no significant differences between the overexpression strains and wild-type worms. Scale bar indicates 25 µm. Error bars indicate SEM. Three biological replicates were performed. Statistical significance was assessed using a one-way ANOVA with Dunnett’s multiple comparisons test. OE = overexpression. ****p<0.0001.

The impact of overexpression of mitochondrial fission or fusion genes on mitochondrial network morphology during aging was also examined, using day 8 adult worms. As we previously reported [17], wild-type worms exhibit increased mitochondrial fragmentation at this aged time point (**Figure 2 C,D**). While worms overexpressing *drp-1, fzo-1* or *eat-3* also exhibit increased mitochondrial fragmentation with age, at the aged time point their mitochondrial morphology was no longer different from wild-type (**Figure 2F-J**).

### Overexpression of mitochondrial fission or fusion genes decreases physiologic rates

To examine the overall health impact of overexpressing mitochondrial fission and fusion genes, general phenotypic traits such as movement, fertility, and development time were evaluated. As motility is frequently used as an indicator of worm health span [43, 44], thrashing rates were quantified for animals at day 1, day 4, and day 8 of adulthood. Overexpression of either *drp-1* or *fzo-1* significantly increased the thrashing rate at all three timepoints, while overexpression of *eat-3* significantly decreased the thrashing rate at all three timepoints (**Figure 3A-C**). Fertility was evaluated by the number of viable progeny per worm. All three overexpression strains had significantly reduced brood sizes (**Figure 3D**). Development times were measured as the time from hatching to adulthood. While overexpression of *drp-1* did not affect development time, overexpression of the mitochondrial fusion genes *fzo-1* and *eat-3* slowed development (**Figure 3E**).

**Figure 3.**
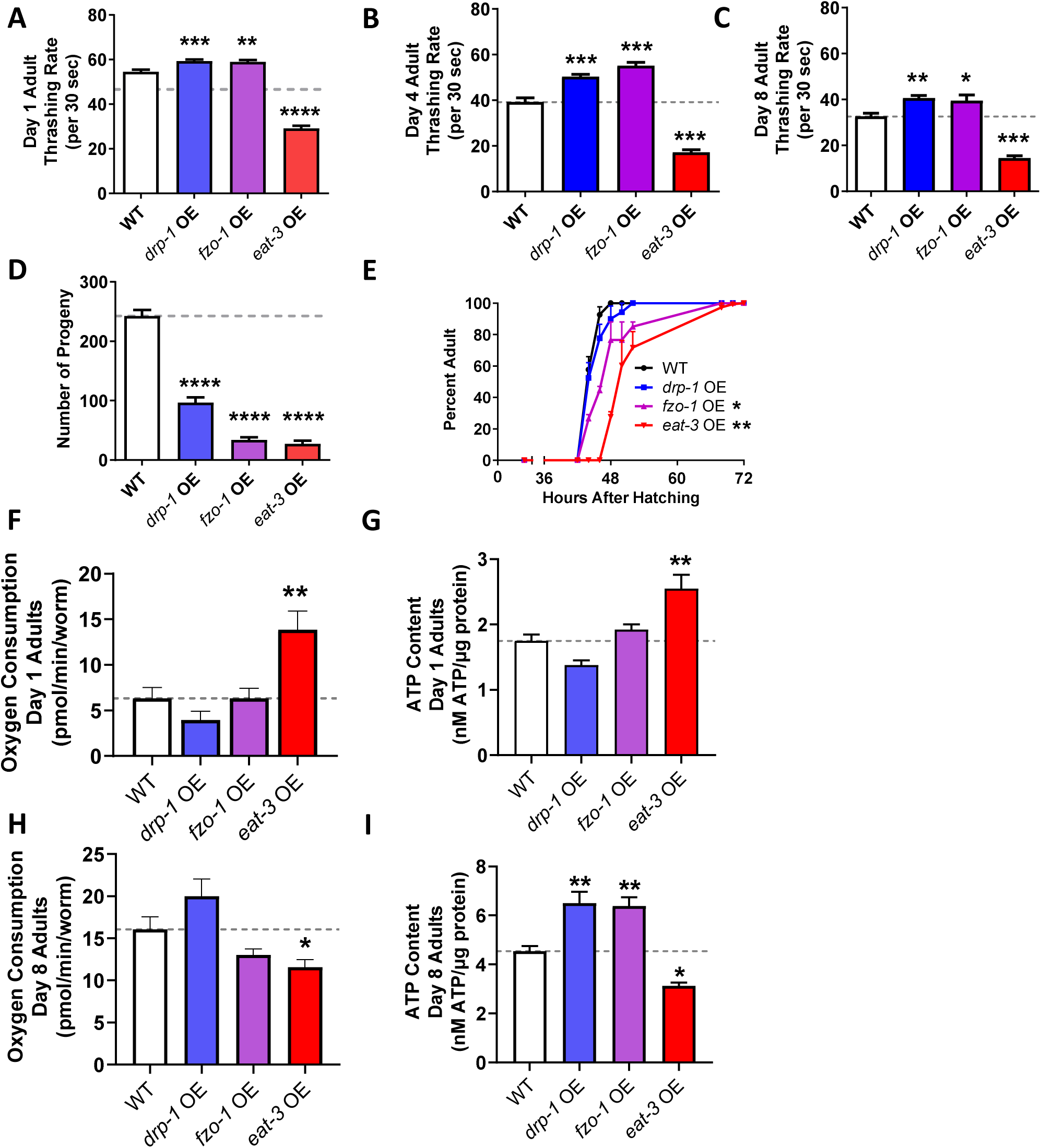
Overexpression of mitochondrial fission or fusion genes results in slowed physiologic rates. The effect of overexpressing mitochondrial fission and fusion genes on the general health of worms was assessed by measuring physiologic rates. (**A-C**) Overexpression of *drp-1* or *fzo-1*resulted in a significant increase in movement as measured by thrashing rate in liquid. In contrast, *eat-3* OE worms exhibited a significant decrease in thrashing rate. (**D**) All three overexpression strains have markedly decreased fertility as indicated by a decreased brood size. (**E**) Overexpression of either of the mitochondrial fusion genes, *fzo-1* or *eat-3*, results in slow post-embryonic development. At day 1 of adulthood, *eat-3* OE worms have increased oxygen consumption (**F**) and increased levels of ATP (**G**). In contrast, at day 8 of adulthood, *eat-3*OE worms have decreased oxygen consumption (**H**) and decreased ATP levels (**I**). Error bars indicate SEM. A minimum of three biological replicates were performed. Statistical significance was assessed using a one-way ANOVA with Dunnett’s multiple comparisons test. OE = overexpression. *p<0.05, **p<0.01, ***p<0.001, ****p<0.0001.

To ensure that the observed phenotypes are specific to overexpression of the mitochondrial fission or fusion genes rather than being an artifact of overexpression in general, we evaluated whether overexpression of an unrelated, control protein could produce similar effects on worm physiology. To do so, we expressed the first exon of the huntingtin protein with a non-disease length polyglutamine tract from the *rpl-28* promoter, which is the same promoter used for the overexpression of *fzo-1*. We found that *rpl-28p::htt19Q* control worms exhibit an increased thrashing rate but a brood size and post-embryonic development time similar to that of wild-type animals (**Figure S1A-C**). This indicates that the deficits in movement, fertility and development that we observed are specific to the overexpression of mitochondrial dynamics genes.

### Overexpression of mitochondrial fission or fusion genes affects mitochondrial function

It has previously been reported that mitochondrial network conformation affects mitochondrial function. Findings from our lab and others suggest that a fragmented mitochondrial network results in a decrease in mitochondrial oxygen consumption and ATP production, both indicators of mitochondrial function [17]. Therefore, we tested whether overexpression of mitochondrial fission or fusion genes would affect mitochondrial function in day 1 and day 8 adults. We found that overexpression of mitochondrial fusion gene *eat-3*, but not *fzo-1*, increased oxygen consumption linked to mitochondrial respiration (**Figure 3F**) and increased ATP content (**Figure 3H**) in day 1 adults. Additionally, we found that overexpression of the mitochondrial fission gene *drp-1* decreased oxygen consumption linked to mitochondrial respiration (**Figure 3F**) as well as ATP content (**Figure 3H**) in day 1 adults, though not to a statistically significant extent. By contrast, at day 8 of adulthood, overexpression of *eat-3* significantly decreased both oxygen consumption linked to mitochondrial respiration as well as ATP content, while overexpression of *drp-1* and *fzo-1* both had increased ATP levels but without significant differences in in oxygen consumption (**Figure 3G, 3I**).

### Overexpression of mitochondrial fusion genes increases resistance to exogenous stressors

Mitochondrial network morphology changes in response to environment, including conditions of stress, in order to adapt to the needs of the cell. Mitochondrial fragmentation can facilitate clearing of damaged mitochondria by mitophagy, while mitochondrial fusion can promote complementation of mitochondrial components and increased mitochondrial function. We therefore examined whether overexpression of mitochondrial fission or fusion genes impacted organismal resistance to heat stress (37°C), osmotic stress (600 mM NaCl), acute oxidative stress (300 µM juglone), chronic oxidative stress (4 mM paraquat), anoxic stress (72 hours, 24 hours recovery) and bacterial pathogen stress (*P. aeruginosa* strain PA14).

Overexpression of the mitochondrial fusion genes *eat-3* and *fzo-1* significantly increased resistance to all six exogenous stressors that we tested (**Figure 4A-F**). Overexpression of the mitochondrial fission gene *drp-1* also provided increased resistance to all the exogenous stressors except for heat stress, where a trends towards increased resistance was observed but failed to reach significance (**Figure 4A-F**). For all stress assays, *eat-3* OE worms exhibited the greatest resistance to stress, followed by *fzo-1* OE worms and then *drp-1* OE worms.

**Figure 4.**
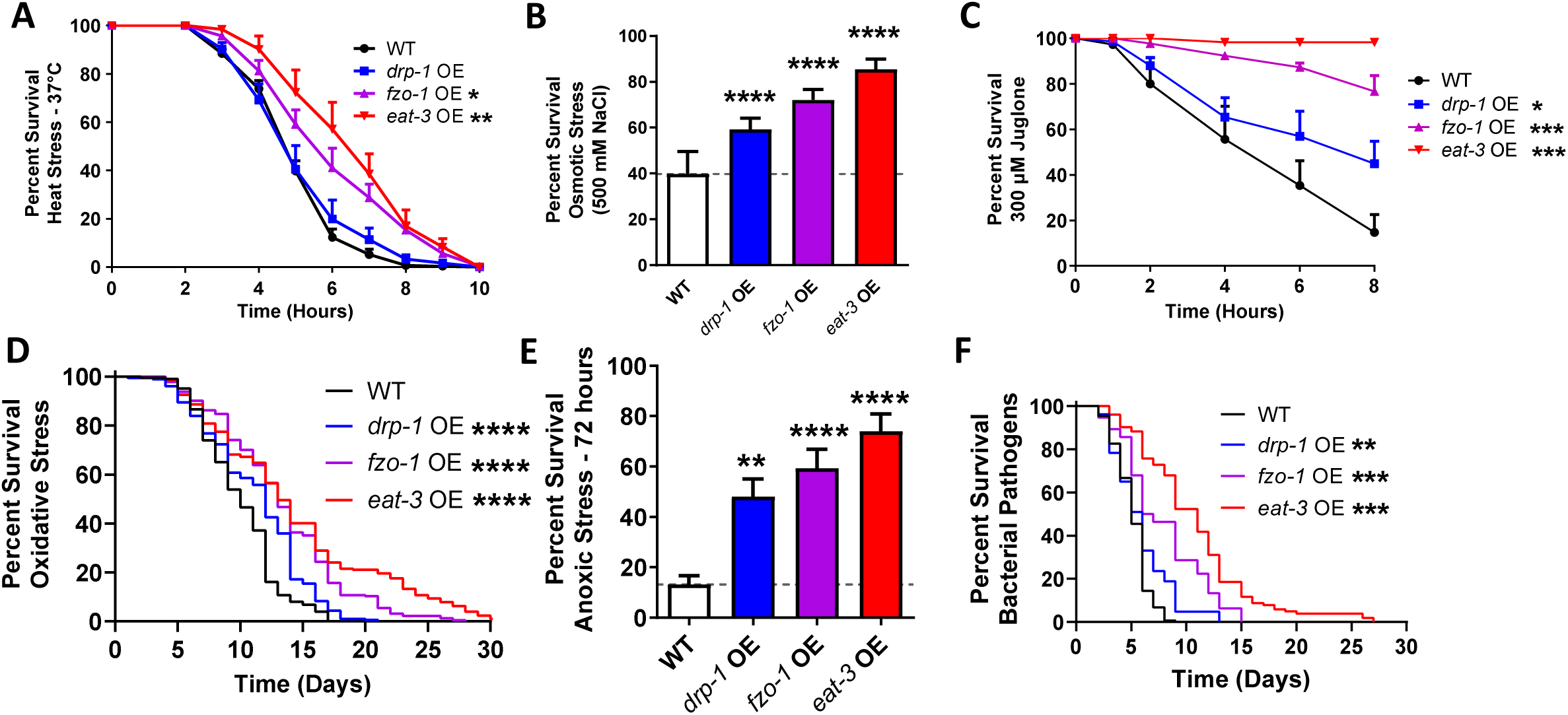
Overexpression of mitochondrial fission or fusion genes increases resistance to exogenous stressors. Overexpression of the mitochondrial fusion genes, *fzo-1* or *eat-3*, results in increased resistance to heat stress at 37°C (**A**). *drp-1* OE, *fzo-1* OE and *eat-3* OE worms all have increased resistance to osmotic stress (500 mM NaCl, **B**), acute oxidative stress (300 µM juglone, **C**), chronic oxidative stress (4 mM paraquat, **D**), anoxia (72 hours, **E**), and bacterial pathogen stress (*P. aeruginosa* strain PA14, **F**). Error bars indicates SEM. A minimum of six biological replicates were performed. Statistical significance was assessed using a repeated measures ANOVA with Tukey’s multiple comparison test in panels A and C, a log-rank test in panels B, D and F, and a one-way ANOVA with Dunnett’s multiple comparison test in panel E. OE = overexpression. *p<0.05, **p<0.01, ***p<0.001, ****p<0.0001.

The *rpl-28p::htt19Q* control survived similarly to wild-type animals in response to heat stress, osmotic stress, chronic oxidative stress and anoxic stress (**Figure S1**). In response to acute oxidative stress, *rpl-28p::htt19Q* worms have increased survival compared to wild-type animals and performed similarly to *drp-1* OE and *fzo-1* OE worms (**Figure S1F**). In response to bacterial pathogen stress, *rpl-28p::htt19Q* worms again had increased survival compared to wild-type worms and performed similarly to *drp-1* OE animals (**Figure S1I**).

### Overexpression of mitochondrial fission or fusion genes extends lifespan

To evaluate whether overexpression of mitochondrial fission or fusion genes could benefit *C. elegans* longevity, we measured the lifespan of animals overexpressing mitochondrial fusion genes *eat-3* and *fzo-1* as well as animals overexpressing the mitochondrial fission gene *drp-1*. Somewhat unexpectedly, we found that overexpression of *drp-1* significantly increases lifespan (**Figure 5A**). Additionally, overexpression of *eat-3* and *fzo-1* nearly doubled wild-type lifespan (**Figure 5B,C**). In contrast, the *rpl-28p::htt19Q* control strain only had a slight increase in lifespan compared to wild-type animals, and lived significantly shorter than animals overexpressing *drp-1, fzo-1* or *eat-3* (**Figure S1J,K**). This indicates that the lifespan extension we observed is specific to the overexpression of mitochondrial fission or fusion genes and not due to overexpression of any gene.

**Figure 5.**
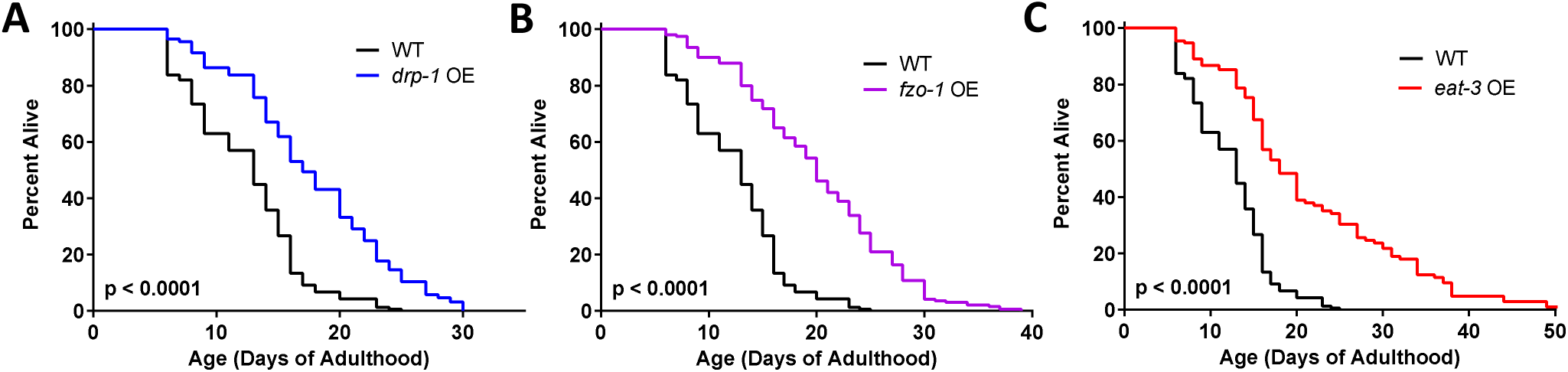
Overexpression of mitochondrial fission or fusion genes extends lifespan. Overexpression of the mitochondrial fission gene *drp-1*or either of the mitochondrial fusion genes *fzo-1* or *eat-3* significantly increases lifespan. Six biological replicates were performed. Statistical significance was assessed using a log-rank test.

### Disruption of mitochondrial fission or fusion genes can extend lifespan

Having shown that overexpression of mitochondrial fission and fusion genes extends longevity, we next examined how disruption of mitochondrial fission and fusion genes affects lifespan. While disruption of *fzo-1* did not affect lifespan, disruption of either *drp-1* or *eat-3* significantly extended lifespan, though to a much lesser extent than overexpression of these genes (**Figure 6A-C**). Additionally, we observed that *fis-1;fis-2* and *mff-1;mff-2* double mutants had reduced lifespans while their single mutant counterparts had no change in lifespan (**Figure 6D-I**), suggesting the possibility of functional redundancy.

**Figure 6.**
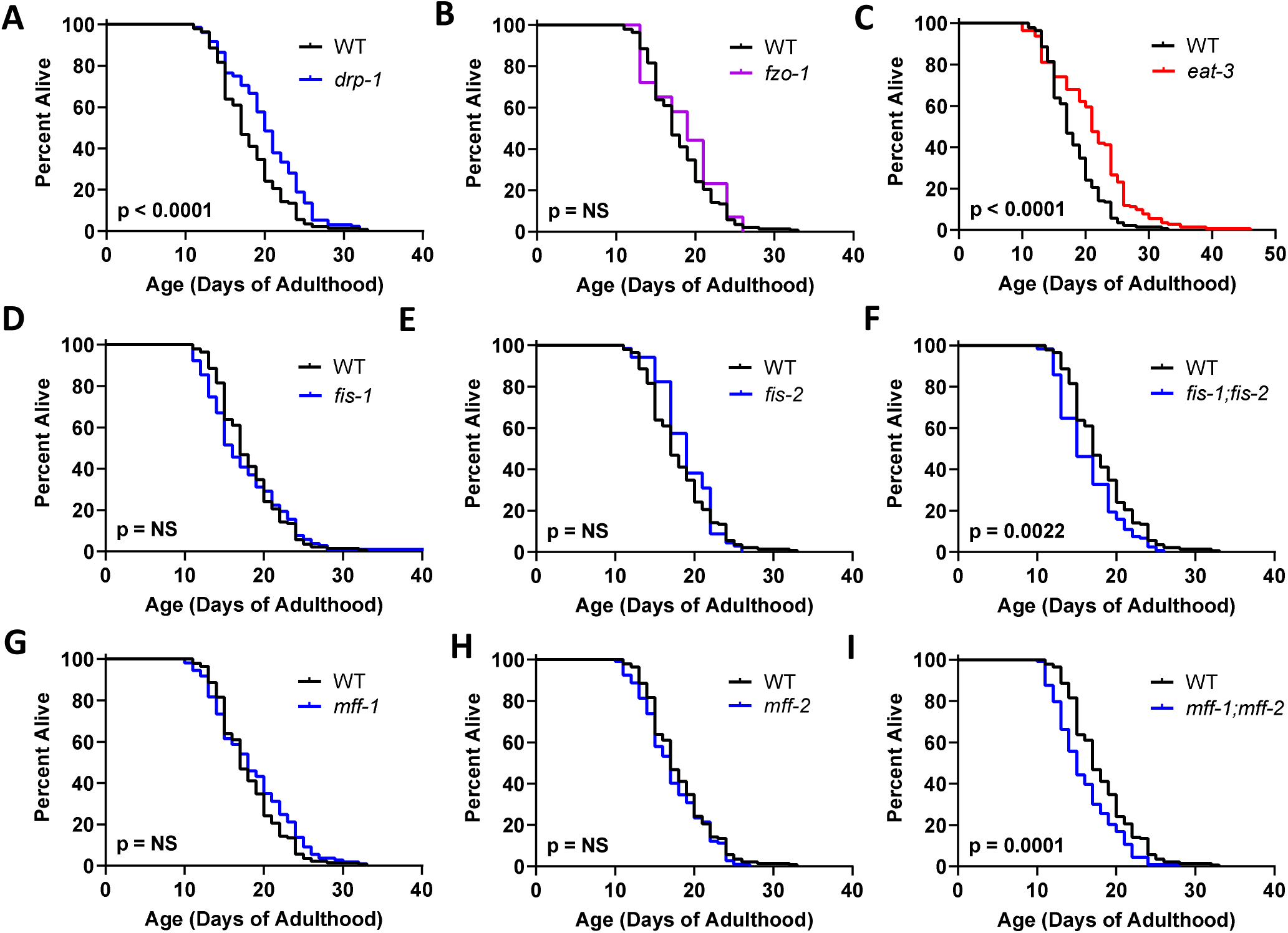
Disruption of mitochondrial fission or fusion genes can extend lifespan. (**A**) Disruption of the mitochondrial fission gene *drp-1* resulted in a small increase in lifespan. While deletion of the mitochondrial fusion gene *fzo-1* did not affect lifespan (**B**), disruption of *eat-3* extended longevity (**C**). Disruption of either *fis-1* (**D**), or *fis-2* (**E**) individually did not affect lifespan, while *fis-1;fis-2* double mutants exhibited a slight decrease in longevity (**F**). Similarly, *mff-1* (**G**) and *mff-2* (**H**) single mutants have a normal lifespan, while *mff-1;mff-2* double mutants exhibit decreased longevity (**I**). At least three biological replicates were performed. Statistical significance was assessed using a log-rank test.

### Overexpression of mitochondrial fusion genes activates multiple pathways of cellular resilience

Activation of key pathways of cellular resilience such as the DAF-16-mediated stress response, the p38-mediated innate immune signaling pathway, the mitochondrial unfolded protein response, the cytosolic unfolded protein response, the SKN-1-mediated oxidative stress response and the HIF-1-mediated hypoxia response can enhance resistance to stress and contribute to lifespan extension [35, 36, 45–48]. To determine if activation of cellular resilience pathways contributes to the lifespan extension and stress resistance in the mitochondrial fission and fusion overexpression strains, we measured mRNA levels of target genes for each pathway using quantitative RT-PCR.

We examined target genes from the mitochondrial unfolded protein response (*cdr-2, F15B9.10, hsp-6;* **Figure 7A-C**), the cytoplasmic unfolded protein response (*hsp-16.2;* **Figure 7D**), the ER-unfolded protein response (*hsp-4*; **Figure 7E**), the hypoxia response (*nhr-57, F22B5.4*; **Figure 7F,G**), the DAF-16-mediated stress response (*mtl-1, sod-3, dod-3, sodh-1*; **Figure 7H-K**), the p38-mediated innate immune signaling pathway (*sysm-1, Y9C9A.8*; **Figure 7L,M**) and the SKN-1-mediated oxidative stress response (*gst-4*; **Figure 7N**). Overexpression of the mitochondrial fusion gene *eat-3* resulted in upregulated expression of multiple DAF-16 target genes, including *mtl-1, sod-3* and *dod-3* (**Figure 7H-J**), as well as target genes from the innate immune signaling pathway (**Figure 7L**) and the SKN-1-mediated oxidative stress response (**Figure 7N**).

**Figure 7.**
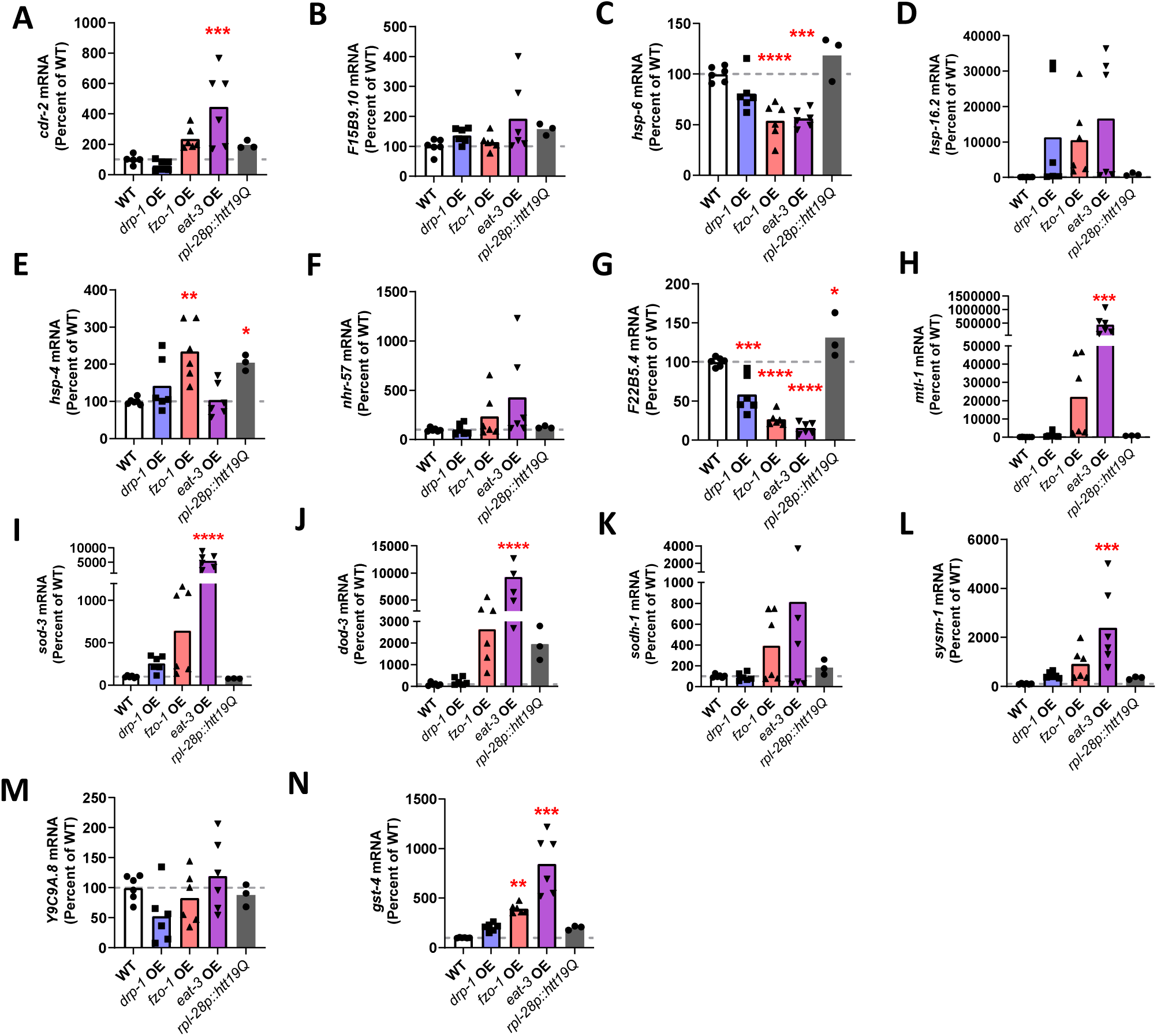
Overexpression of mitochondrial fusion genes activates multiple pathways of cellular resilience. Quantitative RT-PCR was used to assess the expression of target genes from different pathways of cellular resilience in the mitochondrial fission and fusion overexpression strains (*drp-1* OE, *fzo-1* OE and *eat-3* OE). As a control, expression was also examined in worms overexpressing a control protein under the *rpl-28*promoter, which was used in the *fzo-1* OE strain. We examined target genes from the mitochondrial unfolded protein response (*cdr-2, F15B9.10, hsp-6;***A-C**), the cytoplasmic unfolded protein response (*hsp-16.2;* **D**), the ER-unfolded protein response (*hsp-4*; **E**), the hypoxia response (*nhr-57, F22B4.5*; **F,G**), the DAF-16-mediated stress response (*mtl-1, sod-3, dod-3, sodh-1*; **H-K**), the p38-mediated innate immune signaling pathway (*sysm-1, Y9C9A.8*; **L,M**) and the SKN-1-mediated oxidative stress response (*gst-4*; **N**). At least one of the mitochondrial fusion mutants exhibited a significant increase in expression of targets of the DAF-16-mediated stress response pathway (*mtl-1, sod-3, dod-3*), the innate immune signaling pathway *(sysm-1*) and the SKN-1-mediated oxidative stress response *(gst-4*), which was not observed in the control strain. This indicates that multiple pathways of cellular resilience are activated in mitochondrial fusion overexpression strains. Six biological replicates were performed. Statistical significance was assessed using a one-way ANOVA with Dunnett’s multiple comparisons test. *p<0.05, **p<0.01, ***p<0.001, ****p<0.0001.

Overexpression of the mitochondrial fusion gene *fzo-1* also exhibited a trend towards increased expression of these target genes but it only reached significance for the SKN-1 pathway.

Overexpression of *drp-1* did not significantly increase the expression of any of the target genes. With the exception of *hsp-4* and *F22B5.4*, the *rpl-28p::htt19Q* control strain did not increase expression of any target genes, indicating that overexpression of the mitochondrial fission or fusion proteins specifically activated these stress response pathways.

### Overexpression of *drp-1* does not ameliorate mitochondrial morphology or enhance increased stress resistance and lifespan caused by overexpression of mitochondrial fusion genes

As *drp-1* and *fzo-1/eat-3* perform opposite functions within the cell, we hypothesized that overexpression of *drp-1* could restore mitochondrial network morphology in *fzo-1* OE and *eat-3* OE worms, and revert the phenotypic differences observed in these worms to those of wild-type worms. Accordingly, we crossed *drp-1* OE worms to both *fzo-1* OE and *eat-3* OE worms to generate *fzo-1* OE;*drp-1* OE and *eat-3* OE;*drp-1* OE double transgenic worms.

At day 1 of adulthood, *drp-1* OE failed to revert mitochondrial morphology towards wild-type morphology in either *fzo-1* OE or *eat-3* OE animals (**Figure S2A,B**). In fact, *drp-1* OE further decreased mitochondrial area and length and further increased circularity in *eat-3* OE worms. Similarly, at day 8 of adulthood, overexpression of *drp-1* decreased mitochondrial area and length and increased circularity in both *fzo-1* OE and *eat-3* OE worms (**Figure S2A,C**).

In examining the effect of *drp-1* OE on physiologic rates, we found that it was able to restore the thrashing rate of *eat-3* OE worms towards wild-type levels (**Figure S2D**) but did not improve either fertility (**Figure S2E**) or development time (**Figure S2F**). Despite increasing resistance to stress in wild-type worms, overexpression of *drp-1* in *eat-3* OE worms decreased resistance to heat stress (**Figure S3A**), acute oxidative stress (**Figure S3C**), chronic oxidative stress (**Figure S3D**) and bacterial pathogen stress but did not significantly reduce resistance to osmotic stress (**Figure S3B**) or anoxia (**Figure S3E**). *drp-1* OE also significantly decreased the lifespan of both *fzo-1* OE and *eat-3* OE worms (**Figure S3G,H**), despite increasing wild-type lifespan.

### Overexpression of *fzo-1* and *drp-1* in *eat-3* OE worms decreases stress resistance and lifespan

As FZO-1 fuses the outer mitochondrial membrane and EAT-3 fuses the inner mitochondrial membrane, it may be necessary to increase the expression of both genes to increase mitochondrial fusion. To determine if some of the phenotypes we observed might result from an imbalance in fusion of the inner and outer mitochondrial membrane, we characterized *eat-3* OE;*fzo-1* OE double mutants. We also examined *eat-3* OE;*fzo-1* OE;*drp-1* OE triple mutants, which have increased expression of all three of the major mitochondrial fission and fusion genes, and might be predicted to have increased mitochondrial fission and fusion capacity. Similar to overexpression of *eat-3* or *fzo-1* individually, *eat-3* OE;*fzo-1* OE double mutants and *eat-3* OE;*fzo-1* OE;*drp-1* OE triple mutants exhibited decreased thrashing (**Figure S4A**), decreased fertility (**Figure S4B**), and slow post-embryonic development time (**Figure S4C**). Resistance to heat stress (**Figure S4D**), acute oxidative stress (**Figure S4E**), chronic oxidative stress (**Figure S4F**), bacterial pathogens (**Figure S4G**), osmotic stress (**Figure S4H**) and anoxia (**Figure S4I**) in *eat-3* OE;*fzo-1* OE and *eat-3* OE;*fzo-1* OE;*drp-1* OE worms was all significantly diminished compared to *eat-3* OE worms. Similarly, while *eat-3* OE;*fzo-1* OE double mutants and *eat-3* OE;*fzo-1* OE;*drp-1* OE triple mutants live longer than wild-type worms, their lifespan is significantly decreased compared to *eat-3* OE worms (**Figure S4J**). Overall, the overexpression of *fzo-1* or of both *fzo-1* and *drp-1* decreases resistance to stress and lifespan in *eat-3* OE worms.

## Discussion

### Overexpression of mitochondrial fusion genes can cause mitochondrial fragmentation

Contrary to our prediction that overexpression of mitochondrial fusion genes would result in a more fused mitochondrial network, we found that overexpression of either mitochondrial fission or fusion genes both caused mitochondrial fragmentation. However, other groups have also observed mitochondrial fragmentation in response to overexpression of mitochondrial fusion genes. In mammalian cells, the overexpression of the outer mitochondrial fusion proteins MFN1 and MFN2 causes either clustering of spherical mitochondria or elongated mitochondria [49, 50]. Overexpression of the mammalian inner mitochondrial fusion protein OPA1 increases mitochondrial network fragmentation, likely due to an imbalance in the generation of long and short OPA-1 isoforms, the latter of which has been suggested to promote fission [16, 51–55]. Additionally, modest overexpression of OPA1 induces mitochondrial fusion and mitochondrial network elongation while high levels of OPA1 overexpression induces mitochondrial fragmentation, indicating that outcomes from fusion machinery overexpression may be dependent upon the degree of overexpression [56].

In *C. elegans*, the overexpression of either of the mitochondrial fusion genes *fzo-1* or *eat-3* using a heat-shock inducible promoter produces mitochondrial fragmentation in embryos and it was suggested that, as with OPA1, EAT-3 may contribute to fission activity in certain conditions [57]. In the same study, it was reported that overexpression of the outer mitochondrial fusion gene *fzo-1* specifically in the muscle results in mitochondrial fragmentation in the muscle, indicating that overexpression of either of the mitochondrial fusion genes can induce mitochondrial fragmentation in *C. elegans*.

### Overexpression of mitochondrial fusion genes activates multiple stress response pathways

We found that overexpression of the mitochondrial fusion genes *fzo-1* and *eat-3* increases resistance to all tested exogenous stressors and activated multiple stress response pathways. Overexpression of *drp-1* also increases resistance to all stressors other than heat stress, though to a lesser degree compared to *eat-3* OE and *fzo-1* OE. However, we did not observe an increase in the activation of stress response pathways in *drp-1* OE animals, suggesting that altering the expression of *drp-1* does not induce resistance to stress in the same manner as overexpression of the mitochondrial fusion genes.

### Overexpression of mitochondrial fission or mitochondrial fusion genes increases lifespan

Others previously reported that ubiquitous overexpression of the mitochondrial fusion gene *fzo-1* is not sufficient to extend lifespan in *C. elegans* [33]. However, tissue-specific overexpression of *fzo-1* in either the neurons, the muscle or the intestine of animals with a *fzo-1* null mutant background resulted in a slight but significant lifespan extension, suggesting that specific conditions may be required in order for overexpression of *fzo-1* to increase longevity [33].

We predicted that increasing the capacity for mitochondrial fusion events to occur by increasing mitochondrial fusion gene expression could benefit lifespan by decreasing age-associated mitochondrial fragmentation and improving mitochondrial function. We found that, despite increasing mitochondrial fragmentation, ubiquitous overexpression of the mitochondrial fission gene *drp-1* or either of the mitochondrial fusion genes *eat-3* and *fzo-1* significantly extends *C. elegans* lifespan. Thus, our findings suggest that lifespan extension by overexpression of mitochondrial fusion genes may not be dependent on mitochondrial network morphology or function, but rather on activation of stress response pathways triggered by excess levels of mitochondrial fission and fusion proteins. Furthermore, our findings suggest that increasing mitochondrial fragmentation by increasing expression of *drp-1* can extend lifespan and somewhat improve mitochondrial function during aging, contrary to our prediction that decreasing mitochondrial fragmentation benefits longevity.

### Increasing or decreasing expression of mitochondrial fission or fusion genes extends lifespan and increases resistance to stress

Previous reports as to how the deletion of mitochondrial fission and fusion genes affects *C. elegans* lifespan differ [33, 34, 42, 58]. We find that in the case of *eat-3* and *drp-1*, either deletion or overexpression increases lifespan, though deletion of these genes extends lifespan to a much lesser extend compared to overexpression (**Table S1**). Previously, we have also shown that disruption of mitochondrial fusion genes *fzo-1* and *eat-3* increases resistance to stress and activates multiple stress response pathways, similarly to overexpression of mitochondrial fusion genes. We have also observed that while disruption of *drp-1* increases resistance to stress, similarly to overexpression of *drp-1*, it does not activate multiple stress response pathways[17].

We previously reported that disruption of *drp-1* does not affect mitochondrial morphology and that disruption of *eat-3* causes mitochondrial fragmentation. Here, we find that overexpression of either *drp-1* or *eat-3* induces segmentation of the mitochondrial network. Combined, these findings suggest that lifespan extension by increasing or decreasing mitochondrial fission and fusion gene expression is not dependent on either mitochondrial fragmentation or mitochondrial elongation.

Furthermore, while disruption of *drp-1* somewhat decreases mitochondrial function and disruption of *eat-3* significantly decreases mitochondrial function, overexpression of *drp-1* provides an age-associated improvement in mitochondrial function and overexpression of *eat-3* results in an age-associated decline in mitochondrial function [17]. Together, this suggests that lifespan extension by manipulation of mitochondrial fission and fusion gene expression is not specifically linked to the increased or decreased mitochondrial function that may be expected with morphological changes.

Altogether, these findings suggest that an imbalanced expression level of mitochondrial fission or fusion genes may lead to a hormetic response that acts independent of mitochondrial structure or function, leading to increased lifespan possibly through increased expression of pro-survival genes. However, our findings suggest that while increasing or decreasing mitochondrial fusion gene expression benefits lifespan by activating stress response pathways, increasing or decreasing fission gene expression may act to extend lifespan through an alternative mechanism. Given that disruption of *drp-1* significantly extends the lifespan of already long-lived mutants such as *daf-2*, and we have shown that overexpression of *drp-1* significantly extends the lifespan of wild-type animals, future investigations should determine how overexpression of *drp-1* may affect the lifespan of *daf-2* worms and other long-lived mutants [41].

### Simultaneous overexpression of mitochondrial fission and fusion genes is not more beneficial than overexpression of either a fission or a fusion gene

We predicted that the simultaneous overexpression of both *drp-1* and either *fzo-1* or *eat-3* would maintain the dynamicity of mitochondria that is lost with age, improving an organism’s ability to respond to stress, and thus allowing for an increase in lifespan. However, we found that while *fzo-1;drp-1* OE animals still survived longer than wild-type animals, their lifespans were shorter than *fzo-1* OE animals and their mitochondrial networks were more fragmented than wild-type animals. Likewise, the lifespans of *eat-3;drp-1* OE animals were shorter than *eat-3* OE animals and like *eat-3* OE worms, their mitochondrial networks were highly fragmented. Similarly, while *fzo-1;drp-1* OE and *eat-3;drp-1* OE animals still had increased resistance to multiple stressors, they did not survive as well as animals that overexpressed only one gene, suggesting that a more unbalanced expression of mitochondrial fission and fusion genes may promote a stronger stress response despite producing similar mitochondrial morphologies.

We also evaluated whether the benefits of overexpressing both mitochondrial fusion genes *fzo-1* and *eat-3* could further extend lifespan and stress resistance. We again found that the overexpression of both genes resulted in reduced resistance to stress and reduced lifespan extension compared to overexpression of only one fusion gene. The overexpression of all three mitochondrial dynamics genes performed similarly to the overexpression of only the fusion genes, further indicating that a stronger imbalance in the expression of mitochondrial fission or fusion genes may yield a stronger activation of stress response genes and improved survival.

## Conclusion

Overall, our findings demonstrate that overexpression of mitochondrial fission genes or mitochondrial fusion genes both result in mitochondrial fragmentation, increased resilience and extended longevity. We find that as with the deletion of mitochondrial fusion genes, the overexpression of mitochondrial fusion genes activates multiple stress response pathways, which likely contributes to their enhanced resistance to multiple stressors. Combined with our previous work, this indicates that increasing or decreasing the expression of genes involved in mitochondrial fission or mitochondrial fusion can lead to increased stress resistance and lifespan.

## Experimental Procedures

### Strains

WT/N2

JVR587 *Peft-3::drp-1::unc-54 3’UTR*

JVR588 *Ppro-1-eat-3-unc-54 3’UTR*

JVR589 *Prpl-28-fzo-1-unc-54 3’UTR*

JVR617 *drp-1 o/e[Peft-3::drp-1+Pmyo-3::mCherry]; fzo-1 o/e[Prpl-28::fzo-1+Pmyo-2::mCherry]*

JVR618 *drp-1 o/e[Peft-3::drp-1+Pmyo-3::mCherry]; eat-3 o/e[Ppro-1::eat-3+Pvha-6::mCherry]*

JVR122 *bcIs78(pMyo-3::mitoGFP(matrixGFP) + pRF4); rol-6(su1006)II*

JVR622 *drp-1 o/e sybIs3765[Peft-3::drp-1+Pmyo-3::mCherry]; bcIs78(pmyo-3::mitoGFP (matrix GFP) + pRF4); rol-6(su1006)II*

JVR623 *fzo-1 o/e sybIs3776[Prpl-28::fzo-1+Pmyo-2::mCherry]; bcIs78(pmyo-3::mitoGFP (matrix GFP) + pRF4); rol-6(su1006)II*

JVR624 *eat-3 o/e [Ppro-1::eat-3+Pvha-6::mCherry]; bcIs78(pmyo-3::mitoGFP (matrix GFP) + pRF4); rol-6(su1006)II*

JVR625 *drp-1 o/e[Peft-3::drp-1+Pmyo-3::mCherry]; fzo-1 o/e[Prpl-28::fzo-1+Pmyo-2::mCherry]; bcIs78(pmyo-3::mitoGFP (matrix GFP) + pRF4); rol-6(su1006)II*

JVR626 *drp-1 o/e[Peft-3::drp-1+Pmyo-3::mCherry]; eat-3 o/e[Ppro-1::eat-3+Pvha-6::mCherry]; bcIs78(pmyo-3::mitoGFP (matrix GFP) + pRF4); rol-6(su1006)II*

JVR642 *fzo-1 o/e sybIs3776[Prpl-28::fzo-1+Pmyo-2::mCherry]; eat-3 o/e [Ppro-1::eat-3+Pvha-6::mCherry]; drp-1 o/e sybIs3765[Peft-3::drp-1+Pmyo-3::mCherry]*

JVR643 *fzo-1 o/e sybIs3776[Prpl-28::fzo-1+Pmyo-2::mCherry]; eat-3 o/e [Ppro-1::eat-3+Pvha-6::mCherry]*

sybIs6440 *Prpl-28::HTT(Q19)::wrmScarlet::unc-54 3’UTR*

### Quantitative Real-Time RT-PCR

To perform quantitative RT-PCR, we first collected worms in M9 buffer and extracted RNA using Trizol as previously described [59]. Using a High-Capacity cDNA Reverse Transcription kit (Applied Biosystems 4368814), the collected mRNA was then converted to cDNA. Quantitative PCR was performed using a PowerUp SYBR Green Master Mix (Applied Biosystems A25742) in a MicroAmp Optical 96-Well Reaction Plate (Applied Biosystems N8010560) and a Viia 7 Applied Biosystems qPCR machine. mRNA levels were calculated as the copy number of the gene of interest relative to the copy number of the endogenous control, *act-3*, then expressed as a percentage of wild-type. Primer sequences for each target gene are as follows: *drp-1* (L-GAGATGTCGCTATTATCGAACG, R-CTTTCGGCACACTATCCTG) *fzo-1* (L-GCTTTCTGCAGGTTGAAGGT, R-CGACACCAGGGCTATCAAGT) *eat-3* (L-GCGAAGTTTTGGACTTGCTC, R-CGATCGAATCCGAACTGTTT).

### Confocal Imaging and Quantification

Mitochondrial morphology was imaged using worms that express mitochondrially-targeted GFP in the body wall muscle cells as well as a *rol-6* mutant background. The *rol-6* mutation results in animals moving in a twisting motion, allowing one side of the sheaths of muscle cells to always be facing the objective lens and thus facilitating imaging of mitochondrial networks within cells. Without the *rol-6* mutation, only the longitudinal edges of the muscle will often be visible, thus making it difficult to observe mitochondrial organization. Worms at day 1 or day 8 of adulthood were mounted on 2% agar pads and immobilized using 10 µM levamisole. Worms were imaged under a 40× objective lens on a Zeiss LSM 780 confocal microscope. Single plane images were collected for a total of 24 worms over 3 biological replicates for each strain. Imaging conditions were kept the same for all replicates and images. Quantification of mitochondrial morphology was performed using ImageJ. Segmentation analysis was carried out using the SQUASSH (segmentation and quantification of subcellular shapes) plugin. Particle analysis was then used to measure number of mitochondria, mitochondrial area, mitochondrial circularity, and maximum Feret’s diameter (an indicator of particle length).

### Thrashing Rate

Thrashing rates were determined manually by transferring 20 worms onto an unseeded agar plate. One milliliter of M9 buffer was added and the number of body bends per 30 seconds was counted for 3 biological replicates of 6-8 worms per strain.

### Brood Size

Brood size was determined by placing individual prefertile young adult animals onto NGM plates. Worms were transferred to fresh NGM plates daily until progeny production ceased. The resulting progeny was allowed to develop to the L4 stage before quantification. Three biological replicates of 5 animals each were completed.

### Post-Embryonic Development

Postembryonic development (PED) was assessed by transferring ~ 50-100 eggs to agar plates. After 3 h, newly hatched L1 worms were transferred to a new plate. Starting at 28 hours after hatching, worms were scored approximately every 2 hours and worms that reached young adulthood were removed from the plate. The hours from hatching to young adulthood were measured as the PED time. Three biological replicates of 20 animals each were completed.

### Oxygen Consumption Rate

Oxygen consumption measurements were taken using a Seahorse XFe96 analyzer. The night before the assay, probes were hydrated in 200 µL Seahorse calibrant at 37 degrees while the analyzer’s heater was turned off to allow the machine to cool. Day 1 and day 8 worms were collected in M9 buffer and washed three times before being pipetted into a Seahorse 96 well plate (Agilent Technologies Seahorse Flux Pack 103793-100). We pipetted approximately 5-25 worms into each well, a number which others have previously determined to be optimal for this assay [60]. Calibration was performed after 22 µL of FCCP and 24 µL sodium azide was loaded into the drug ports of the sensor cartridge. Measurements began within 30 minutes of worms being added to the wells. Basal oxygen consumption was measured 5 times before the first drug injection. FCCP-induced oxygen consumption was measured 9 times, then sodium-azide induced oxygen consumption was measured 4 times. Measurements were taken over the course of 2 minutes and before each measurement the contents of each well were mixed for an additional 2 minutes. Non-mitochondrial respiration (determined by sodium azide-induced oxygen consumption rate) was subtracted from basal respiration to calculate mitochondrial respiration.

### ATP Determination

Day 1 and day 8 adult worms were collected, washed 3 times and frozen in 50 µL of M9 buffer using liquid nitrogen. Samples were then immersed in boiling water for 15 minutes followed by ice for 5 minutes and finally spun down at 14,800g for 10 minutes at 4 ℃. Supernatants were diluted 10-fold before ATP measurements using a Molecular Probes ATP determination kit (A22066) and TECAN plate reader. Luminescence was normalized to protein content measured using a Pierce BCA protein determination kit.

### Heat Stress Assay

To measure resistance to heat stress, approximately 25 pre-fertile young adult worms were transferred to new NGM plates freshly seeded with OP50 bacteria and were incubated at 37℃. Starting at 2 hours, survival was measured every hour for a total of 10 hours of incubation. Three biological replicates were completed.

### Osmotic Stress Assay

To measure resistance to osmotic stress, approximately 25 pre-fertile young adult worms were transferred to NGM plates containing 500 mM NaCl and seeded with OP50 bacteria. Worms were kept at 20℃ for 24 hours before survival was scored. Five biological replicates were completed.

### Oxidative Stress Assays

Resistance to acute oxidative stress was measured by transferring approximately 25 pre-fertile young adult worms to 300 μM juglone plates seeded with OP50 bacteria. Worms were kept at 20℃ and survival was monitored every 2 hours for a total of 8 hours. Resistance to chronic oxidative stress was performed by transferring 30 pre-fertile young adult worms to freshly prepared plates containing 4 mM paraquat, 25 µM FUdR and seeded with concentrated OP50. Survival was monitored daily. Three biological replicates were completed for both assays.

### Anoxic Stress Assay

To measure resistance to anoxic stress, approximately 50 pre-fertile young adult worms were transferred to new NGM plates seeded with OP50 bacteria. To create a low-oxygen environment for the worms, we utilized Becton-Dickinson Bio-Bag Type A Environmental Chambers. Plates with young adult worms of each strain were placed in the Bio-Bags for 48 hours at 20℃, then removed from the bags and allowed to recover for 24 hours at 20℃ before survival was measured. Five biological replicates were completed.

### Bacterial Pathogen Stress Assay

We tested for nematode resistance to death by bacterial colonization of the intestine. The slow kill assay was performed as previously described [61, 62]. OP50 bacteria was seeded to the center of NGM plates containing 100 mg/L FUdR and plates were left at room temperature for two days. PA14 cultures were grown with aeration at 37℃ for 16 hours, then seeded to the center of NGM agar plates containing 20 mg/L FUdR. The plates seeded with PA14 bacteria were allowed to dry, then incubated at 37℃ for 24 hours and then at room temperature for 24 hours. Approximately 40 L4 worms were transferred to plates containing 100 mg/L FUdR that were seeded with OP50 bacteria, and the worms were grown at 20℃ until they reached day 3 of adulthood. Day 3 adult worms were then transferred from these plates onto plates containing 20 mg/L FUdR that were seeded with PA14 bacteria. The assay was conducted at 20℃ and survival was monitored daily until all worms died. Three biological replicates were completed.

### Lifespan Assay

Lifespan assays were completed at 20℃ and on NGM agar plates that contained FUdR to inhibit the development of progeny and limit internal hatching. We used a low concentration of 25 μM FUdR, which we have previously shown does not affect the longevity of wild-type worms [63]. For each lifespan assay, 40 pre-fertile young adult worms were transferred to 25 μM FUdR plates seeded with OP50 bacteria and were kept at 20℃. Four biological replicates were started on four subsequent days and all replicates were scored every other day to monitor survival until all worms died. Worms were excluded from the assay if they crawled off the agar and died on the side of the plate, had internal hatching of progeny or expulsion of internal organs. Raw lifespan data are provided in **Table S2**.

### Statistical Analysis

A minimum of three biological replicates were completed for all assays. Where possible, the experimenter was blinded to the genotype during the course of the experiment, to ensure unbiased results. Statistical significance of differences between groups was determined by computing a t-test, a one-way ANOVA, a two-way ANOVA or a log-rank test using Graphpad Prism, as indicated in the Figure legends. All error bars indicate the standard error of the mean.

## SUPPLEMENTAL FIGURES

**Figure S1.**
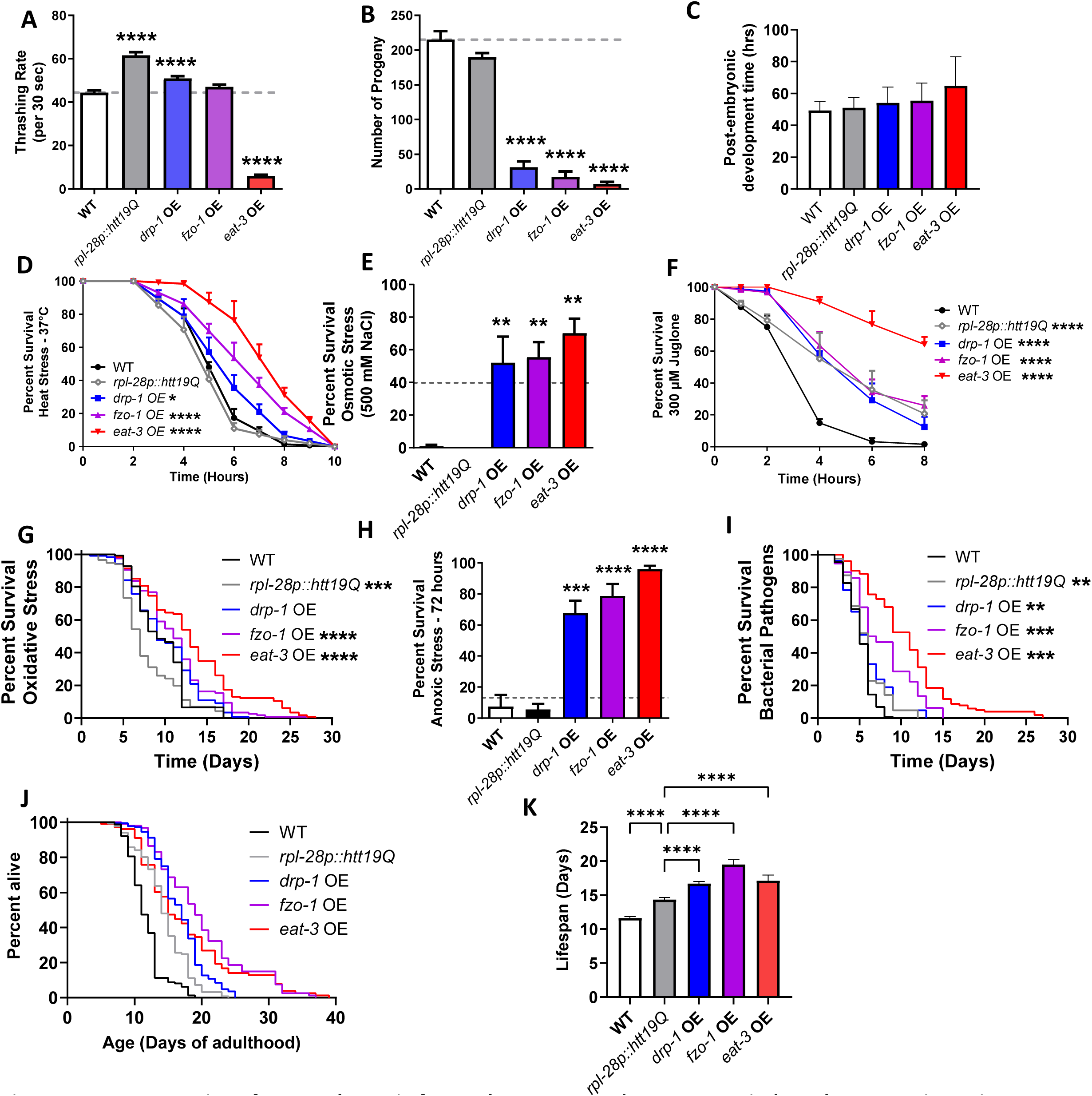
Overexpression of a control protein from *rpl-28* promoter does not recapitulate phenotypes in strains overexpressing mitochondrial fission or fusion genes. To control for overexpression, we compared the phenotype of *drp-1* OE, *fzo-1*OE and *eat-3*OE worms to worms overexpressing a control protein (exon 1 of wild-type huntingtin) under the *rpl-28* promoter that was used to generate *fzo-1*OE worms. *rpl-28p::htt19Q*worms did not have decreased movement (**A**) or decreased fertility (**B**). These worms also exhibited wild-type post-embryonic development (**C**) as well as resistance to heat stress (**D**) and osmotic stress (**E**). *rpl-28p::htt19Q* did show a significantly increased resistance to acute oxidative stress (**F**) suggesting the possibility that this phenotype might not be specific to overexpression of mitochondrial fission and fusion genes. Expression of *htt19Q* from the *rpl-28* promoter did not increase resistance to oxidative stress (G) or anoxia (H). While *rpl-28::htt19Q* worms have a small increase in resistance to bacterial pathogens (**I**) and lifespan (**J,K**), this increase is less than observed in the mitochondrial fusion overexpression strains. Error bars indicate SEM. A minimum of six biological replicates were performed. Statistical significance was assessed using a one-way ANOVA with Dunnett’s multiple comparison test in A,B, C, E, H and K; a repeat measures ANOVA with Tukey’s multiple comparison test in D and F; and a log-rank test in G, I and J. OE = overexpression. Data on overexpression strains from Figures 3,4 and 5. *p<0.05, **p<0.01, ***p<0.001, ****p<0.0001.

**Figure S2.**
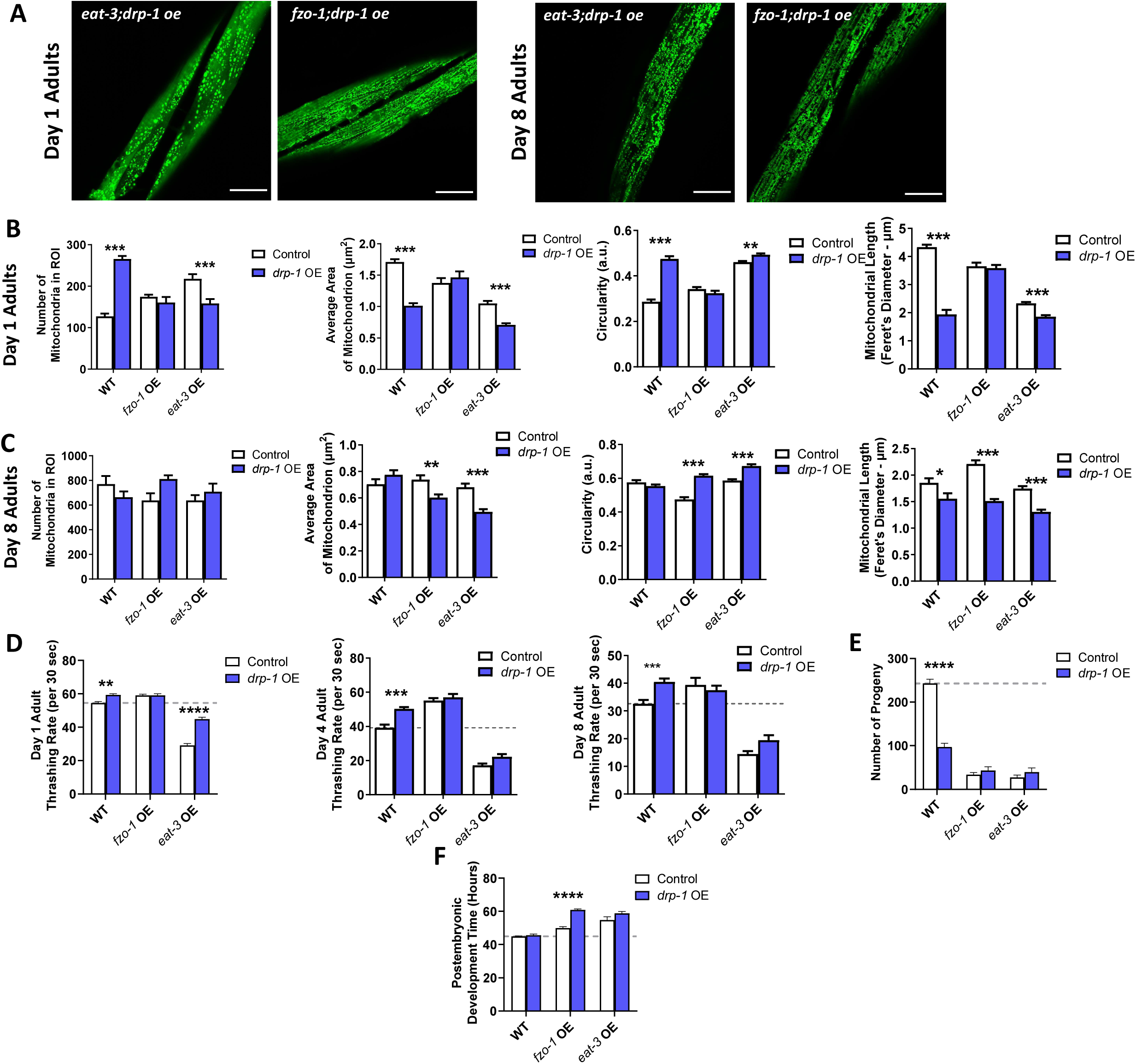
Overexpression *drp-1* causes further mitochondrial fragmentation in *eat-3* OE worms. To determine if overexpression of the mitochondrial fission *drp-1* would diminish phenotypes caused by overexpression of mitochondrial fusion genes, *drp-1* OE worms were crossed with *fzo-1* OE and *eat-3* OE worms. (**A**) Images of mitochondrial morphology in *eat-3;drp-1* and *fzo-1;drp-1* worms at day 1 and day 8 of adulthood. Scale bar indicates 25 µm. (**B**) At day 1 of adulthood, overexpression of *drp-1* decreased mitochondrial number, decreased mitochondrial area, increased mitochondrial circularity and decreased mitochondrial length in *eat-3* OE worms. *drp-1* OE did not affect the mitochondrial morphology of *fzo-1* OE worms. (**C**) At day 8 of adulthood, overexpression of *drp-1* decreased mitochondrial area, increase mitochondrial circularity and decreased mitochondrial length in both *fzo-1* OE and *eat-3* OE worms.(**D**) *drp-1* OE ameliorated the decreased movement of *eat-3* OE worms on day 1 of adulthood but not at day 4 or day 8. (**E**) Overexpression of *drp-1* did not affect fertility in *fzo-1* OE or *eat-3* OE worms and resulted in a slowing of post-embryonic development time in *fzo-1* OE worms (**F**). Errors bars indicate SEM. A minimum of three biological replicates were performed. Statistical significance was assessed using a two-way ANOVA with Šidák’s multiple comparisons test. OE = overexpression. *p<0.05, **p<0.01, ***p<0.001, ****p<0.0001.

**Figure S3.**
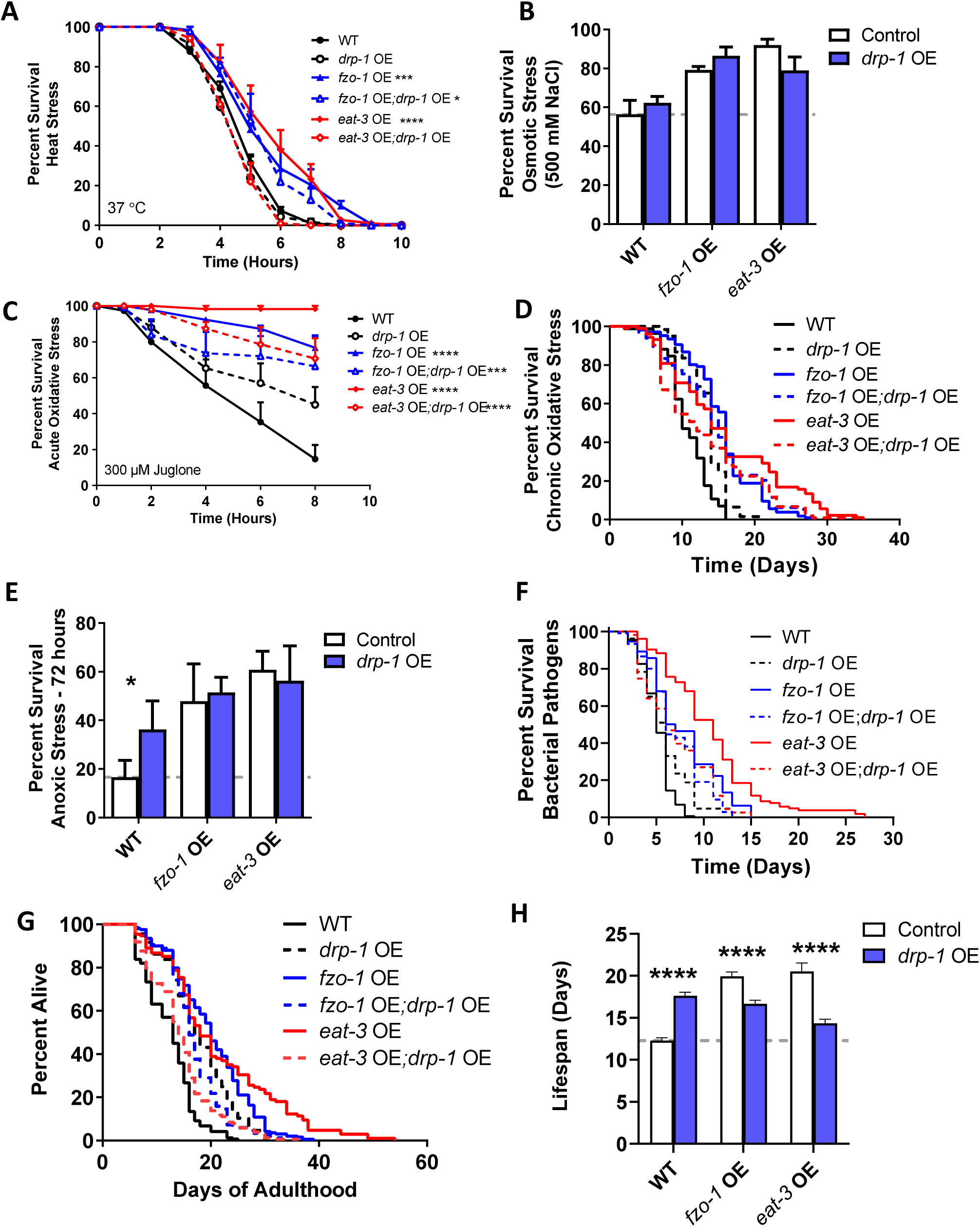
Overexpression *drp-1* decreases stress resistance and lifespan in worms overexpressing *eat-3.* (**A**) Overexpression of *drp-1* reverts heat stress resistance to wild-type in *eat-3* OE worms. (**B**) *eat-3* OE;*drp-1* OE worms exhibit a trend toward decreased osmotic stress resistance compared to *eat-3* OE worms. (**C**) *drp-1* OE decreases acute oxidative stress resistance in *fzo-1* OE and *eat-3* OE worms. (**D**) *drp-1* OE also reduces resistance to chronic oxidative stress in *eat-3* OE worms. (**E**) Overexpression of *drp-1* does not affect anoxia resistance in *fzo-1* OE or *eat-3* OE worms. While *drp-1* OE increases bacterial pathogen resistance (**F**) and lifespan (**G,H**) in wild-type worms, it decreases resistance to bacterial pathogens and lifespan in worms overexpressing mitochondrial fusion genes. Error bars indicate SEM. A minimum of three biological replicates were performed. Statistical significance was assessed using a repeated measures ANOVA with Tukey’s multiple comparison test in A and C; a two-way ANOVA with Šidák’s multiple comparisons test in B, E, and H; and a log-rank test in D, F and G. OE = overexpression. Data on single overexpression strains from Figures 4 and 5. *p<0.05, **p<0.01, ***p<0.001, ****p<0.0001.

**Figure S4.**
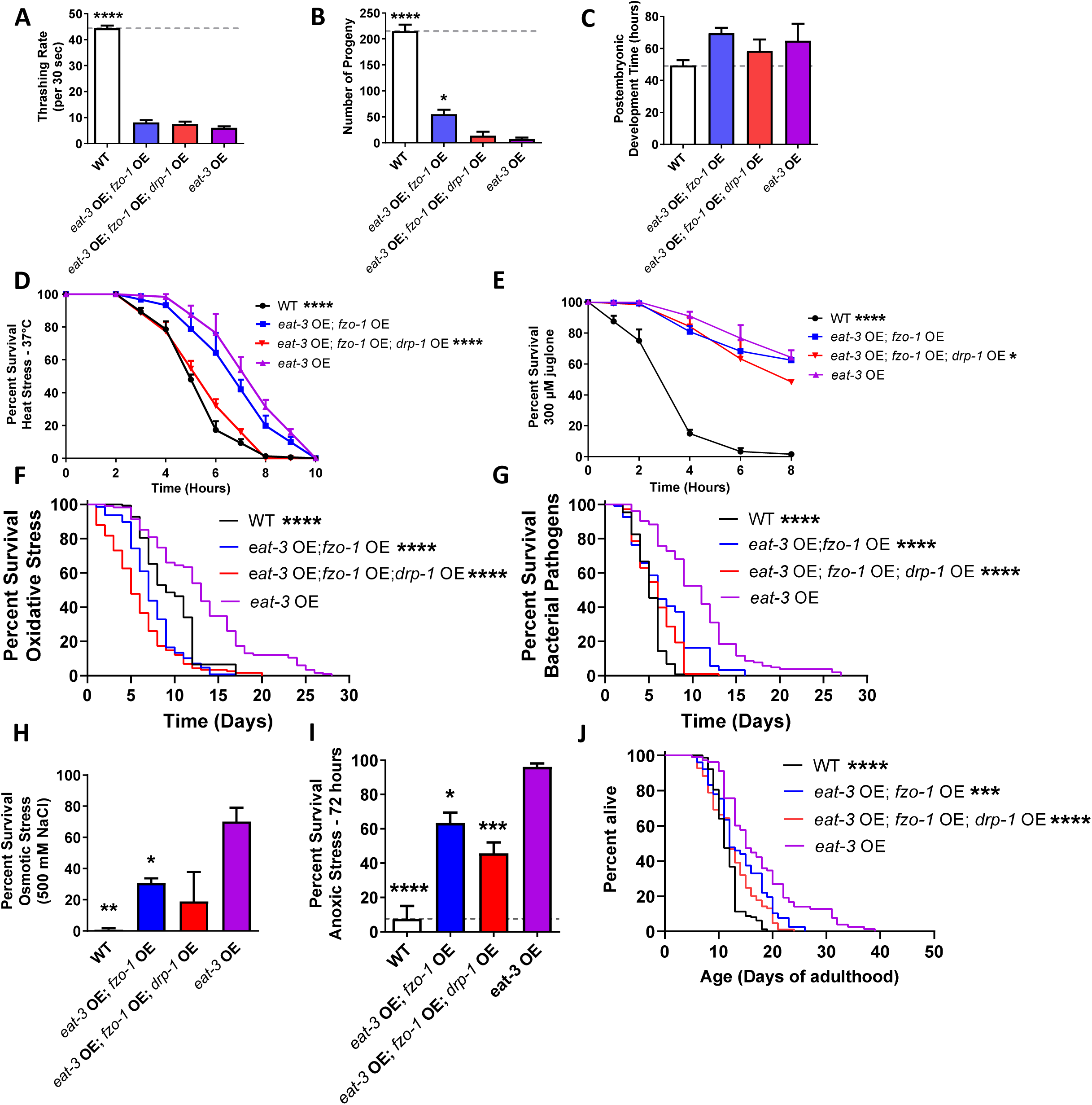
Effects of overexpressing *drp-1, fzo-1* and *eat-3* are not additive. To examine the effects of increasing the expression of *drp-1, fzo-1* and *eat-3*simultaneously, we generated *eat-3* OE; *fzo-1* OE; *drp-1* OE worms. Similar to *eat-3* OE worms, *eat-3* OE; *fzo-1*OE; *drp-1* OE worms exhibit decreased movement (**A**), reduced fertility (**B**), and slow post-embryonic development (**C**). These worms have only a small increase in resistance to heat stress (D). While *eat-3* OE; *fzo-1* OE; *drp-1* OE worms have increased resistance to acute oxidative stress (E), they have increased sensitivity to chronic oxidative stress (F). *eat-3*OE; *fzo-1* OE; *drp-1* OE worms have a small increase in resistance to bacterial pathogens, which is less than observed in *eat-3* OE worms (G). These worms also exhibit a small increase in resistance to osmotic stress (H) and anoxia (I). Although *eat-3* OE; *fzo-1* OE; *drp-1* OE worms have increased lifespan compared to wild-type worms, the magnitude of increase is much less than in *eat-3*worms. Error bars indicate SEM. A minimum of three biological replicates were performed. Statistically significant differences from *eat-3* OE worms are indicated. Statistical significance was assessed using a one-way ANOVA with Dunnett’s multiple comparison test in A, B, C, H and I; a repeated measures ANOVA with Tukey’s multiple comparison test in D and E; and a log-rank test in F, G and J. OE = overexpression. *p<0.05, **p<0.01, ***p<0.001, ****p<0.0001.

**Table S1.** Summary of phenotypes observed in strains with modulated expression of mitochondrial fission and fusion genes.

|  | *drp-1* OE | *fzo-1* OE | *eat-3* OE | *drp-1* DEL | *fzo-1* DEL | *eat-3* DEL |
| --- | --- | --- | --- | --- | --- | --- |
| Mito number | ↑ | ↑ | ↑ | = | ↑ | ↑ |
| Mito. size | ↓ | ↓ | ↓ | = | ↓ | ↓ |
| Circularity | ↑ | ↑ | ↑ | = | ↑ | ↑ |
| Feret's diameter | ↓ | ↓ | ↓ | = | ↓ | ↓ |
| ATP levels (day 1) | = | = | ↑ | ↓ | ↓ | ↓ |
| Oxygen consumption (day 1) | = | = | ↑ | = | ↓ | ↓ |
| Movement (day 1) | ↑ | ↑ | ↓ | ↓ | = | ↓ |
| Fertility | ↓ | ↓ | ↓ | ↓ | ↓ | ↓ |
| Development time | = | ↑ | ↑ | ↑ | ↑ | ↑ |
| Heat stress | = | ↑ | ↑ | ↓ | ↑ | ↑ |
| Osmotic stress | = | = | ↑ | ↓ | ↓ | ↓ |
| Acute oxidative stress | ↑ | ↑ | ↑ | ↑ | ↑ | ↑ |
| Chronic oxidative stress | ↑ | ↑ | ↑ | ↑ | ↑ | ↑ |
| Anoxia | ↑ | = | = | ↓ | ↓ | ↓ |
| Bacterial pathogen stress | ↑ | ↑ | ↑ | ↑ | ↑ | ↑ |
| Lifespan | ↑ | ↑ | ↑ | ↑ | = | ↑ |

